# Selective D_2_ and D_3_ receptor antagonists oppositely modulate cocaine responses in mice via distinct postsynaptic mechanisms in nucleus accumbens

**DOI:** 10.1101/439398

**Authors:** Daniel F. Manvich, Alyssa K. Petko, Rachel C. Branco, Stephanie L. Foster, Kirsten A. Porter-Stransky, Kristen A. Stout, Amy H. Newman, Gary W. Miller, Carlos A. Paladini, David Weinshenker

## Abstract

**Background:** The D_3_ receptor (D_3_R) has emerged as a promising pharmacotherapeutic target for the treatment of several diseases including schizophrenia, Parkinson’s disease, and substance use disorders. However, studies investigating the modulatory impact of D_3_R antagonism on dopamine neurotransmission or the effects drugs of abuse have produced mixed results, in part because D_3_R-targeted compounds often also interact with D_2_ receptors (D_2_R). The purpose of this study was to compare the consequences of selective D_2_R or D_3_R antagonism on the behavioral effects of cocaine in mice, and to identify the neurobiological mechanisms underlying their modulatory effects.

**Methods:** We characterized the effects of selective D_2_R or D_3_R antagonism in mice on 1) basal and cocaine-induced locomotor activity, 2) presynaptic dopamine release and clearance in the nucleus accumbens using *ex vivo* fast scan cyclic voltammetry, and 3) dopamine-mediated signaling in D_1_-expressing and D_2_-expressing medium spiny neurons using *ex vivo* electrophysiology.

**Results:** Pretreatment with the selective D_2_R antagonist L-741,626 attenuated, while pretreatment with the selective D_3_R antagonist PG01037 enhanced, the locomotor-activating effects of acute and repeated cocaine administration. While both antagonists potentiated cocaine-induced increases in presynaptic DA release, D_3_R blockade uniquely facilitated DA-mediated excitation of D_1_-expressing medium spiny neurons in the nucleus accumbens.

**Conclusions:** Selective D_3_R antagonism potentiates the behavioral-stimulant effects of cocaine in mice, an effect that is in direct opposition to that produced by selective D_2_R antagonism or nonselective D_2_-like receptor antagonists, likely by facilitating D_1_-mediated excitation in the nucleus accumbens. These findings provide important insights into the neuropharmacological actions of D_3_R antagonists on mesolimbic dopamine neurotransmission.

## Introduction

Dopamine (DA) dysregulation has been implicated in several neurological and affective disorders including schizophrenia (1, 2), Parkinson’s disease (3, 4), depression (1, 5), and substance use disorders (SUD) (6, 7). Five subtypes of DA receptors have been identified and are grouped into two families: the D_1_-like receptors (D_1_R and D_5_R) and D_2_-like receptors (D_2_R, D_3_R, and D_4_R) (8, 9). D_2_-like receptors serve as the target of numerous clinically-available pharmacotherapeutics for conditions in which disrupted DA signaling has been implicated, most notably Parkinson’s disease (8, 10) and schizophrenia (8, 11). However, undesirable side effects limit the benefits and reduce compliance with such medications (12, 13). These side effects have also contributed to the overall failure of nonselective compounds targeting D_2_-like receptors to show therapeutic potential for SUD (14-16). Consequently, there has been interest in identifying D_2_-like receptor subtype-selective compounds that may engender therapeutic benefits while minimizing aversive side effects (17-19).

The D_3_R exhibits several characteristics that have sparked investigations into its potential as an improved pharmacotherapeutic target for the treatment of DA-related diseases (19-22). Among the D_3_R’s unique properties is a highly restricted pattern of distribution as compared to the D_2_R or D_4_R, with greatest abundance in limbic regions including the nucleus accumbens (NAc) (19, 23-25). Within the NAc, it functions both as a presynaptic inhibitory autoreceptor on DA terminals arising from the ventral tegmental area (8, 26-28) and as a postsynaptic heteroreceptor on GABAergic medium spiny neurons (MSNs) (25, 29-32). While MSNs can be classified based on expression of either D_1_R (D1-MSNs) or D_2_R (D2-MSNs), both types co-express the D_3_R (30, 33-35), with the D_3_R exhibiting the strongest affinity for DA across all DA receptor subtypes (24, 36). The D_3_R is thus well-suited to influence NAc DA neurotransmission (18, 37). However, studies examining its modulatory impact on NAc DA signaling have produced mixed results. For example, some animal studies indicate that genetic knockout or pharmacological antagonism of the D_3_R may facilitate basal or drug-induced increases in extracellular DA within the NAc (38-44), while others have reported no such effect (38, 41, 43, 45, 46) or even an attenuation of drug-induced DA increases (42). Moreover, studies employing behavioral readouts of NAc DA neurotransmission using unconditioned behavioral responses (e.g. locomotor activity) in rodents have produced a similarly confusing picture. D_3_R knockout or pharmacological antagonism have been reported to increase (47-52), decrease (49, 53, 54), or have no effect (39, 45, 55, 56) on basal or drug-induced increases in locomotion. As none of these studies has identified a uniform underlying mechanism to explain their observed neurochemical or behavioral effects, a clear understanding of how D_3_R antagonism modulates NAc DA neurotransmission and associated behaviors has remained elusive.

The purpose of this study was to definitively determine the impact of selective D_3_R antagonism on NAc DA neurotransmission when administered either alone or under conditions of enhanced DA tone, achieved via administration of the monoamine transporter inhibitor and psychostimulant, cocaine. To achieve specific pharmacological D_3_R blockade, we used the highly-selective D_3_R antagonist PG01037, which exhibits ~133-fold selectivity for the D_3_R vs. D_2_R (57, 58). The effects of PG01037 on multiple aspects of NAc DA neurotransmission were assessed and compared to those produced by the selective D_2_R antagonist, L-741,626, which exhibits ~15-42-fold selectivity for the D_2_R vs. D_3_R (59, 60). We first measured changes in basal and cocaine-increased locomotor activity in mice following pretreatment with either of these two selective antagonists and found that they exerted opposing influences on cocaine-induced locomotion. We then addressed potential underlying presynaptic and postsynaptic mechanisms by examining each antagonist’s effect on stimulated presynaptic DA release and DA clearance in the NAc via *ex vivo* fast-scan cyclic voltammetry (FSCV), as well as their impact on the activity of NAc D1-MSNs and D2-MSNs using *ex vivo* whole cell electrophysiology.

## Methods and Materials

### Animals

All procedures were performed in strict accordance with the National Institutes of Health’s Guide for the Care and Use of Laboratory Animals and were approved by the Institutional Animal Care and Use Committees of Emory University or University of Texas at San Antonio. Additional details are provided in the Supplement.

### Effects of Selective D_3_R or D_2_R Antagonism on Locomotor Behavior

The effects of systemic pretreatment with PG01037 or L-741,626 on basal and cocaine-induced locomotor activity were assessed using methods adapted from those previously described (61, 62). Detailed methodological descriptions for drug treatments and locomotor measurements are provided in the Supplement.

### Effects of Selective D_3_R or D_2_R Antagonism on Presynaptic DA Release in NAc

*Ex vivo* slice FSCV was performed as described previously (63, 64) to determine the impact of PG01037 or L-741,626 on stimulated DA release and clearance, and cocaine-induced modulations of these measures, in the NAc. Detailed methods are provided in the Supplement.

### Effects of Selective D_3_R or D_2_R Antagonism on Postsynaptic D1-MSN or D2-MSN Activity in NAc

NAc D1-MSN and D2-MSN activity was assessed using e*x vivo* electrophysiology in slices derived from Drd1a-tdTomato and Drd2-EGFP mice. A detailed description of the recording methods is provided in the Supplement.

### Statistics

Data were analyzed using either paired t-test, one-way repeated measures (RM) analysis of variance (ANOVA), or two-way ANOVA (RM on one or both factors) with post hoc Dunnett’s or Holm-Sidak’s tests as specified. See the Supplement for additional details. GraphPad Prism v. 7.04 (GraphPad Software, La Jolla, CA) was used to perform all statistical analyses. Significance was set at *p* < 0.05 for all tests.

## Results

### Pretreatment with Selective D_2_R or D_3_R Antagonists Produces Opposite Effects on Cocaine-Induced Locomotion

Neither of the active doses of L-741,626 (3.0 or 10.0 mg/kg) significantly affected basal locomotor activity, assessed over the first 30 min following drug administration (Figure 1A). By contrast, cocaine-induced increases in locomotor activity were robustly attenuated by L-741,626 pretreatment (Figure 1B). Analyses revealed that 10.0 mg/kg L-741-626 significantly reduced the locomotor-activating effects of each cocaine dose, shifting the cocaine dose-response function rightward and downward (Figure 1B). The effects of L-741,626 were evident immediately after cocaine administration and persisted throughout the observation period (Supplemental Figure S1A-C).

**Figure 1.**
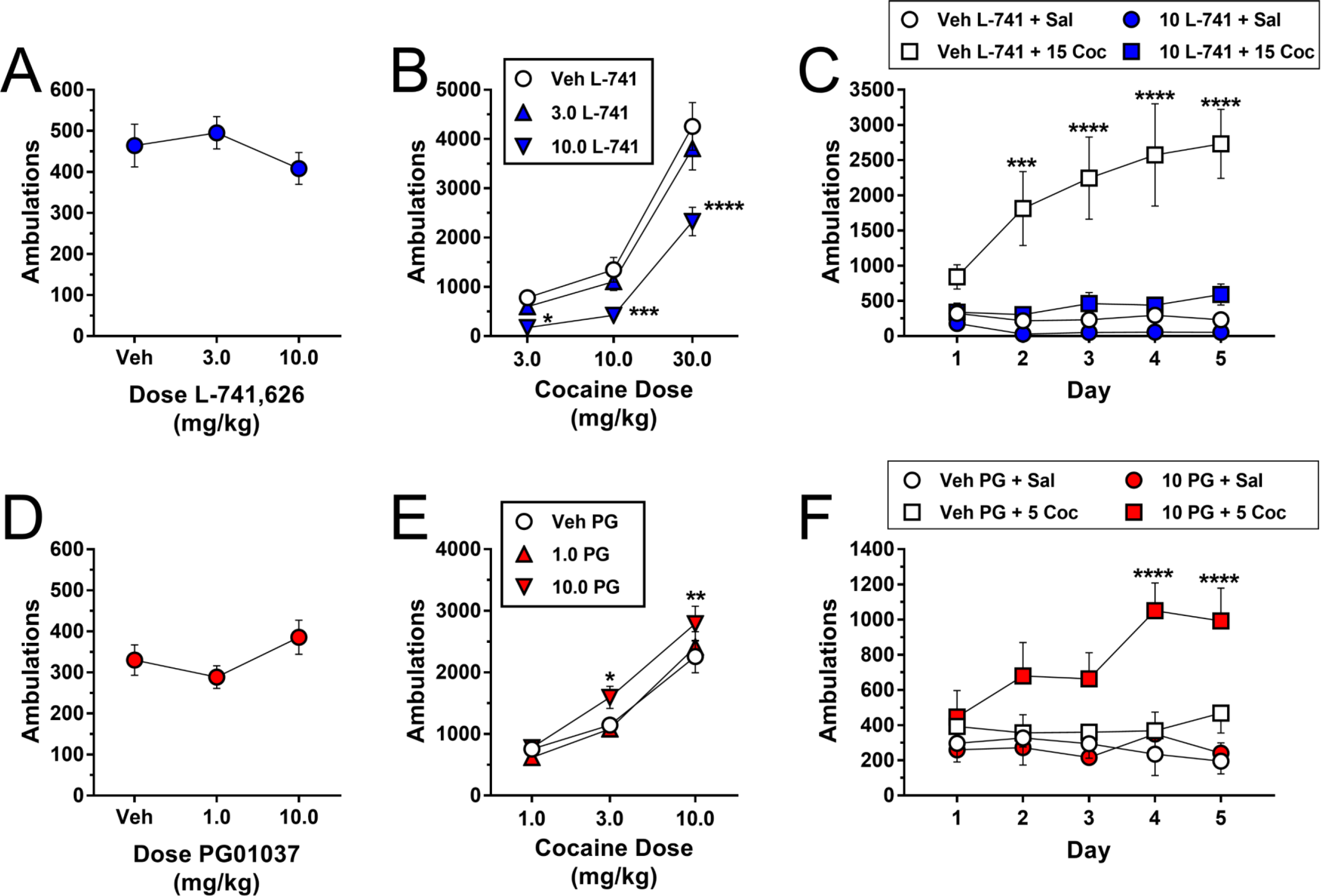
Effects of pretreatment with the D_2_R antagonist L-741,626 or the D_3_R antagonist PG01037 on basal locomotor activity, acute cocaine-induced locomotor activity, and cocaine-induced sensitization in mice. **(A)** Total number of ambulations in the 30-min period following i.p. administration of L-741,626 (one-way RM ANOVA: main effect of dose [*F*_(2,28)_ = 4.90, *p* = 0.015; post hoc Dunnett’s tests comparing active doses to vehicle, *p* > 0.05]). **(B)** Effects of pretreatment with L-741,626 on locomotor activity in the 60-min period following i.p. administration of cocaine (two-way RM ANOVA: main effect of cocaine dose [*F*_(2,28)_ = 82.11, *p* < 0.0001], main effect of L-741,626 dose [*F*_(2,28)_ = 36.17, *p* < 0.0001], interaction [*F*_(4,56)_ = 5.17, *p* = 0.001]). **(C)** Effects of L-741,626 on cocaine-induced sensitization. Mice received one of the following four combinations of L-741,626 and cocaine daily for 5 days: vehicle + saline; vehicle + 15.0 mg/kg cocaine; 10.0 mg/kg L-741,626 + saline; 10.0 mg/kg L-741,626 + 15.0 mg/kg cocaine. Shown are the total number of ambulations in the 60-min period following administration of saline or 15.0 mg/kg cocaine, which was administered 30 min after pretreatment with either 10.0 mg/kg L-741,626 or its vehicle (two-way mixed-design ANOVA: main effect of group [*F*_(3,24)_ = 12.14, *p* < 0.0001], main effect of day [*F*_(4,96)_ = 4.66, *p* = 0.002], interaction [*F*_(12,96)_ = 4.88, *p* < 0.0001]). **(D)** Total number of ambulations in the 30-min period following i.p. administration of PG01037 (one-way RM ANOVA: main effect of dose [*F*_(2,28)_ = 5.87, *p* = 0.007; post hoc Dunnett’s tests comparing active doses to vehicle, *p* > 0.05]). **(E)** Effects of pretreatment with PG01037 on locomotor activity in the 60-min period following i.p. administration of cocaine (two-way RM ANOVA: main effect of cocaine dose [*F*_(2,28)_ = 50.05, *p* < 0.0001], main effect of PG01037 dose [*F*_(2,28)_ = 14.34, *p* < 0.0001], interaction [*F*_(4,56)_ = 1.48, *p* = 0.221]). **(F)** Effects of PG01037 on cocaine-induced sensitization. Mice received one of the following four combinations of PG01037 and cocaine daily for 5 days: vehicle + saline; vehicle + 5.0 mg/kg cocaine; 10.0 mg/kg PG01037 + saline; 10.0 mg/kg PG01037 + 5.0 mg/kg cocaine. Shown are the total number of ambulations in the 60-min period following administration of saline or 5.0 mg/kg cocaine, which was administered 30 min after pretreatment with either 10.0 mg/kg PG01037 or its vehicle (two-way mixed-design ANOVA: main effect of group [*F*_(3,26)_ = 4.95, *p* = 0.008], main effect of day [*F*_(4,104)_ = 2.01, *p* = 0.099], interaction [*F*_(12,104)_ = 2.35, *p* = 0.010]). For panels *A, B, D,* and *E*, all mice (*n* = 15) received all treatments. \**p* < 0.05, \*\**p* < 0.01, \*\*\**p* < 0.001, \*\*\*\**p* < 0.0001, significant difference by Dunnett’s test as compared to vehicle dose of pretreatment drug at the same dose of cocaine. For panels *C* and *F*, mice (*n* = 6-8 per group) received the same drug combination once per day for 5 consecutive days. \*\*\**p* < 0.001, \*\*\*\**p* < 0.0001, significant difference by Dunnett’s test compared to day 1 within the same drug combination group. For all panels, each data point represents mean ± SEM ambulations

Chronic exposure to cocaine produces neurobiological adaptations that cause sensitized behavioral and neurochemical responses to subsequent cocaine challenges (65-67). We wondered whether blockade of D_2_Rs would modulate the induction of locomotor sensitization following repeated exposure to cocaine. Mice treated with the daily combination of vehicle + 15.0 mg/kg cocaine exhibited a robust sensitization of cocaine-induced locomotion, but the development of sensitization was abolished by daily pretreatment with 10.0 mg/kg L-741,626 prior to 15.0 mg/kg cocaine (Figure 1C). Notably, administration of 10.0 mg/kg L-741,626 alone (i.e. prior to saline) did not produce significant changes in locomotor activity (Figure 1C).

We next assessed the effects of pretreatment with the selective D_3_R antagonist PG01037 on acute cocaine-induced increases in locomotion. Neither of the active doses of PG01037 (1.0 or 10.0 mg/kg) significantly affected basal locomotor activity (Figure 1D). In direct opposition to the effects of L-741,626, cocaine-induced increases in locomotor activity were enhanced following pretreatment with PG01037 (Figure 1E). Analyses showed that pretreatment with 10.0 mg/kg PG01037 produced a leftward and upward shift of the cocaine dose-response function (Figure 1E). The potentiation of cocaine-induced locomotion by PG01037 occurred immediately upon cocaine administration and was not attributable to a prolongation of cocaine’s time course of action (Supplemental Figure S2A-C). Based on these results, we hypothesized that PG01037 may similarly potentiate cocaine-induced sensitization. We optimized our sensitization protocol for the detection of enhanced sensitization by testing the effects of PG01037 in conjunction with a low dose of cocaine (5.0 mg/kg) that does not induce behavioral sensitization alone. Mice treated with the daily combination of vehicle + saline, vehicle + 5.0 mg/kg cocaine, or 10.0 mg/kg PG01037 + saline failed to exhibit sensitization. By contrast, sensitized locomotor responses developed in mice pretreated daily with 10.0 mg/kg PG01037 prior to 5.0 mg/kg cocaine (Figure 1F).

Previous work has shown that intra-NAc administration of nonselective D_2_-like antagonists attenuates cocaine-induced locomotor activity (68, 69). However, studies investigating the specific role of the D_3_R subtype in modulating cocaine-induced locomotion have produced mixed results (39, 45, 47-56). We therefore investigated whether intra-NAc infusion of PG01037 would recapitulate the effects of systemic D_3_R blockade described above. In a separate cohort of mice, we delivered bilateral intra-NAc infusions of PG01037 (3 μg/side) or vehicle (aCSF) via guide cannula followed immediately by i.p. administration of 10.0 mg/kg cocaine. Intra-NAc delivery of PG01037 enhanced cocaine-induced locomotor activity (Supplemental Figure S3A-B), suggesting that the NAc is a critical site in which D_3_R antagonists act to potentiate the locomotor-activating effects of cocaine.

### Selective Blockade of D_2_R or D_3_R Facilitates Cocaine-Induced Increases in NAc DA Release

The results described above demonstrate that D_2_R antagonism attenuates, while D_3_R antagonism enhances, the behavioral-stimulant effects of cocaine. Moreover, previous findings (68, 69) and our present work (Supplemental Figure S3) indicated that the NAc is a key neuroanatomical substrate for these effects. However, the neuropharmacological mechanisms underlying these disparate outcomes remained unresolved. Within the NAc, D_2_Rs and D_3_Rs function both as presynaptic autoreceptors on DAergic terminals and as postsynaptic heteroreceptors on MSNs (8, 9), and are thus positioned to modulate presynaptic DA release and postsynaptic DA receptor-mediated changes in neuronal activity, respectively. We therefore sought to address whether D_2_R or D_3_R antagonism similarly or distinctly impact each of these measures.

We first quantified electrically-evoked presynaptic DA release in the NAc in the presence of D_2_R or D_3_R antagonists alone and in combination with cocaine using *ex vivo* slice FSCV. Figures 2A and 2B show representative color plots and current traces for DA detection with increasing concentrations of L-741,626. Application of L-741,626 alone failed to alter peak stimulated DA release (Figure 2C) or the rate of DA clearance (Figure 2D). Figures 2E and 2F show representative color plots and current traces for DA detection following combined application of cocaine (1 μ M) and increasing concentrations of L-741,626. Cocaine enhanced stimulated DA release compared to baseline as reported previously (70, 71), and application of 100 pM L-741,626 potentiated the cocaine-induced increase in DA release (Figure 2G). Application of the highest concentration of L-741,626 (1 nM) trended towards producing this effect as well. Statistical analysis narrowly failed to detect a significant main effect of condition on DA clearance, although cocaine produced an expected increase in tau that was seemingly unaffected by co-application of L-741,626 (Figure 2H).

**Figure 2.**
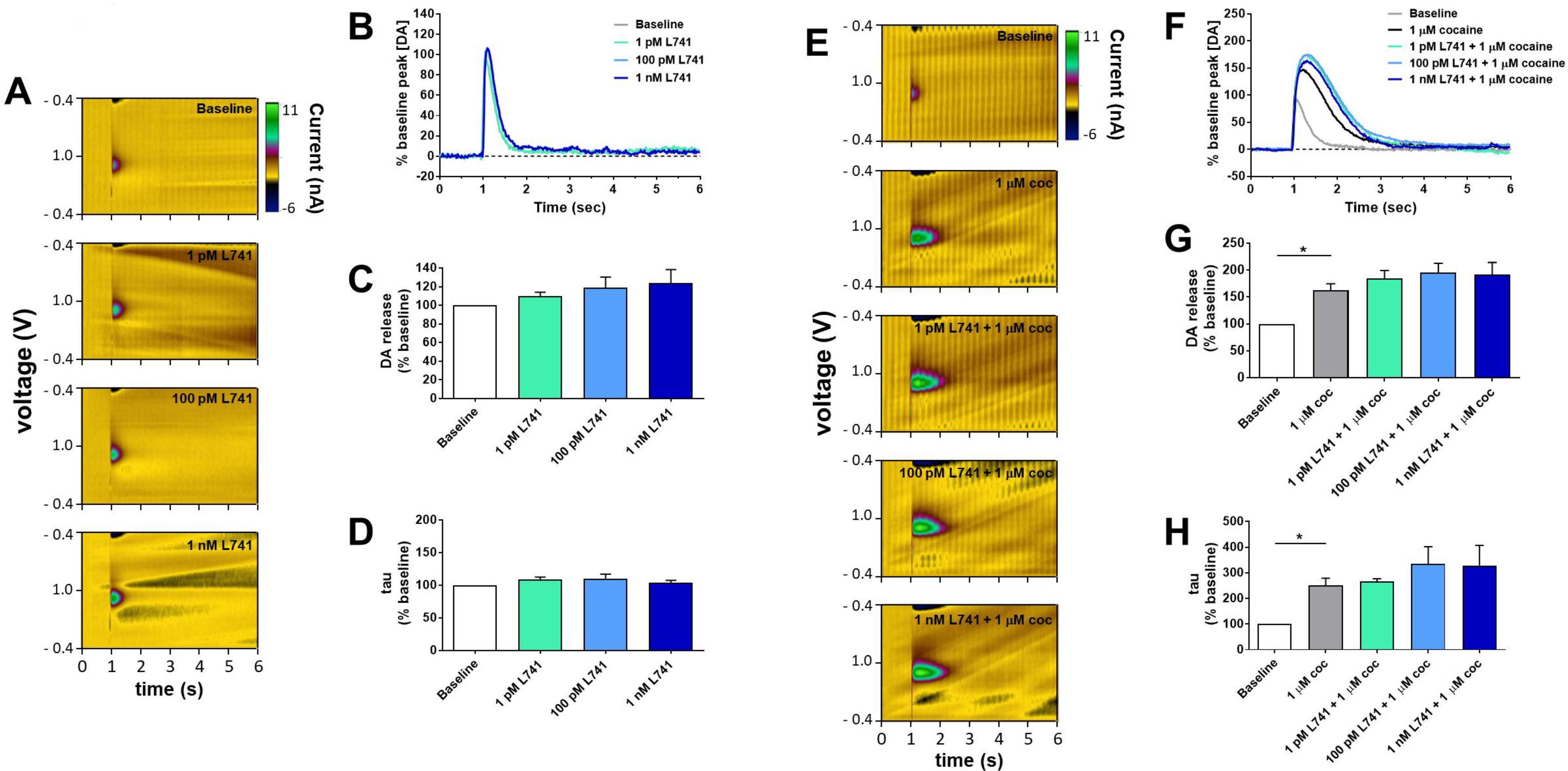
Effects of selective D_2_R antagonism alone or in combination with cocaine on stimulated DA release and DA clearance in the NAc as measured by *ex vivo* fast scan cyclic voltammetry. Representative **(A)** color plots and **(B)** current traces depict electrochemical detection of DA in the presence of varied concentrations of the selective D_2_R antagonist L-741,626. Quantification of **(C)** peak stimulated DA release (one-way RM ANOVA: main effect of concentration [*F*_(3,9)_ = 2.23, *p* = 0.229]) and **(D)** DA clearance (one-way RM ANOVA: main effect of concentration [*F*_(3,9)_ = 1.06, *p* = 0.396]) as a function of applied concentrations of L-741,626 (*n* = 4). Representative **(E)** color plots and **(F)** current traces depict electrochemical detection of DA in the presence of 1 μM cocaine alone and in combination with varied concentrations of L-741,626. Quantification of **(G)** peak stimulated DA release (one-way RM ANOVA: main effect of treatment condition [*F*_(4,16)_ = 18.32, *p* = 0.005]) and **(H)** DA clearance (one-way RM ANOVA: main effect of treatment condition [*F*_(4,12)_ = 6.95, *p* = 0.073]) in the presence of cocaine alone and in combination with L-741,626 (*n* = 5). Color plots in **(A)** and **(E)** depict representative changes in current in z axis (color) with time along the abscissa and applied cyclic potential along the ordinate. Current traces in **(B)** and **(F)** depict release and clearance of DA with time along the abscissa and DA concentration (normalized as a percentage of the mean of all samples collected at the baseline condition) along the ordinate. In **(C)** and **(G)**, values are depicted as the mean ± SEM maximum DA concentration following stimulation (normalized as percentage of the mean peak DA release at baseline). In **(D)** and **(H)**, values are depicted as the mean ± SEM calculated tau constant (normalized as percentage of the mean tau value at baseline) \**p* < 0.05, Dunnett’s test as compared to 1 μM cocaine

Figures 3A and 3B show representative color plots and current traces for DA detection with increasing concentrations of PG01037. Application of PG01037 alone did not alter peak stimulated DA release (Figure 3C) or DA clearance (Figure 3D). Figures 3E and 3F show representative color plots and current traces for DA detection following combined application of cocaine (1 μM) and increasing concentrations of PG01037. Cocaine increased peak stimulated DA release and this effect was further enhanced following application of 100 pM or 1 nM PG01037 (Figure 3G). Cocaine also produced an expected increase in tau value, but in contrast to L-741,626, co-application of PG01037 potentiated this effect (Figure 3H).

**Figure 3.**
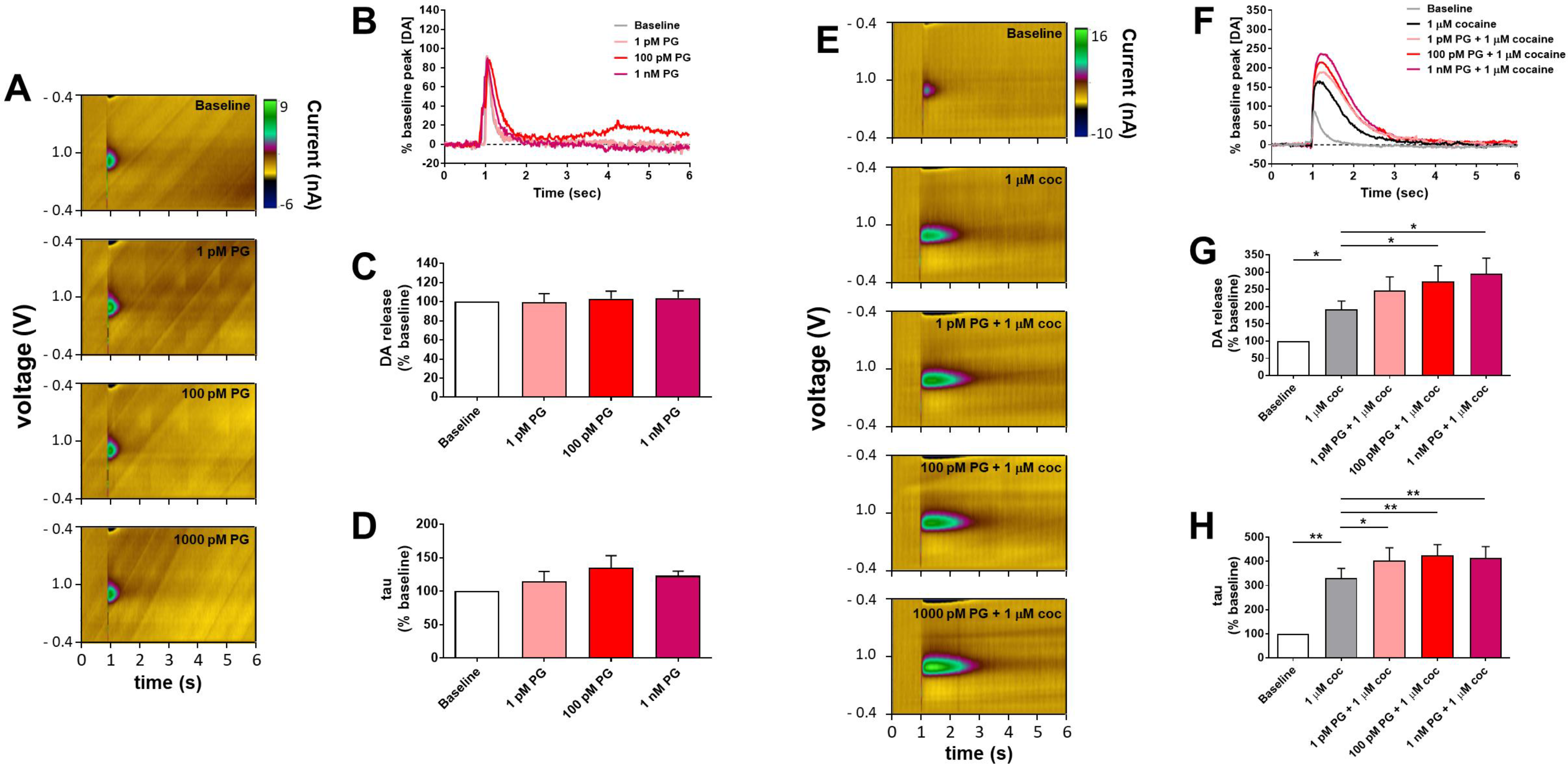
Effects of selective D_3_R antagonism alone or in combination with cocaine on stimulated DA release in the NAc as measured by *ex vivo* fast scan cyclic voltammetry. Representative **(A)** color plots and **(B)** current traces depict electrochemical detection of DA in the presence of varied concentrations of the selective D_3_R antagonist PG01037. Quantification of **(C)** peak stimulated DA release (one-way RM ANOVA: main effect of concentration [*F*_(3,6)_ = 0.18, *p* = 0.837]) and **(D)** DA clearance (one-way RM ANOVA: main effect of concentration [*F*_(3,6)_ = 1.19, *p* = 0.390]) as a function of applied concentrations of PG01037 (*n* = 3). Representative **(E)** color plots and **(F)** current traces depict electrochemical detection of DA in the presence of 1 μM cocaine alone and in combination with varied concentrations of PG01037. Quantification of **(G)** peak stimulated DA release (one-way RM ANOVA: main effect of treatment condition [*F*_(4,24)_ = 13.61, *p* = 0.003]) and **(H)** DA clearance (one-way RM ANOVA: main effect of treatment condition [*F*_(4,24)_ = 34.71, *p* = 0.0001]) in the presence of cocaine alone and in combination with PG01037 (*n* = 7). \**p* < 0.05, \*\**p* < 0.01, Dunnett’s test as compared to 1 μM cocaine. Additional details (e.g. units of measurement, axes, etc.) are as described in Figure 2.

### Selective Blockade of D_2_R or D_3_R Produces Differential Effects on Neuronal Firing in NAc D1-MSNs, but not D2-MSNs

We next examined potential postsynaptic differences by performing *ex vivo* whole cell electrophysiology to examine changes in activity of NAc D1-MSNs and D2-MSNs following combined application of DA and selective D_2_R or D_3_R antagonists. Shown in Figure 4A-D are the effects of L-741,626 on DA-mediated changes in D1-MSN activity. DA application alone significantly increased spike frequency as compared to baseline, but co-administration of L-741,626 did not modulate this effect of DA (Figure 4B). Neither the application of DA alone nor the co-application of DA + L-741,626 affected F-I slope (Figure 4C), and the DA-mediated reduction of rheobase in D1-MSNs was similarly unaffected following co-administration with L-741,626 (Figure 4D). By contrast, application of PG01037 modulated several DA-mediated changes in D1-MSN activity (Figure 4E-H). Combined administration of DA + PG01037 increased spike frequency as compared to both baseline and DA alone conditions, an observation that was accompanied by an increase in the slope of the F-I curve (Figure 4G). Furthermore, application of DA significantly reduced rheobase as compared to baseline, and PG01037 potentiated this effect (Figure 4H).

**Figure 4.**
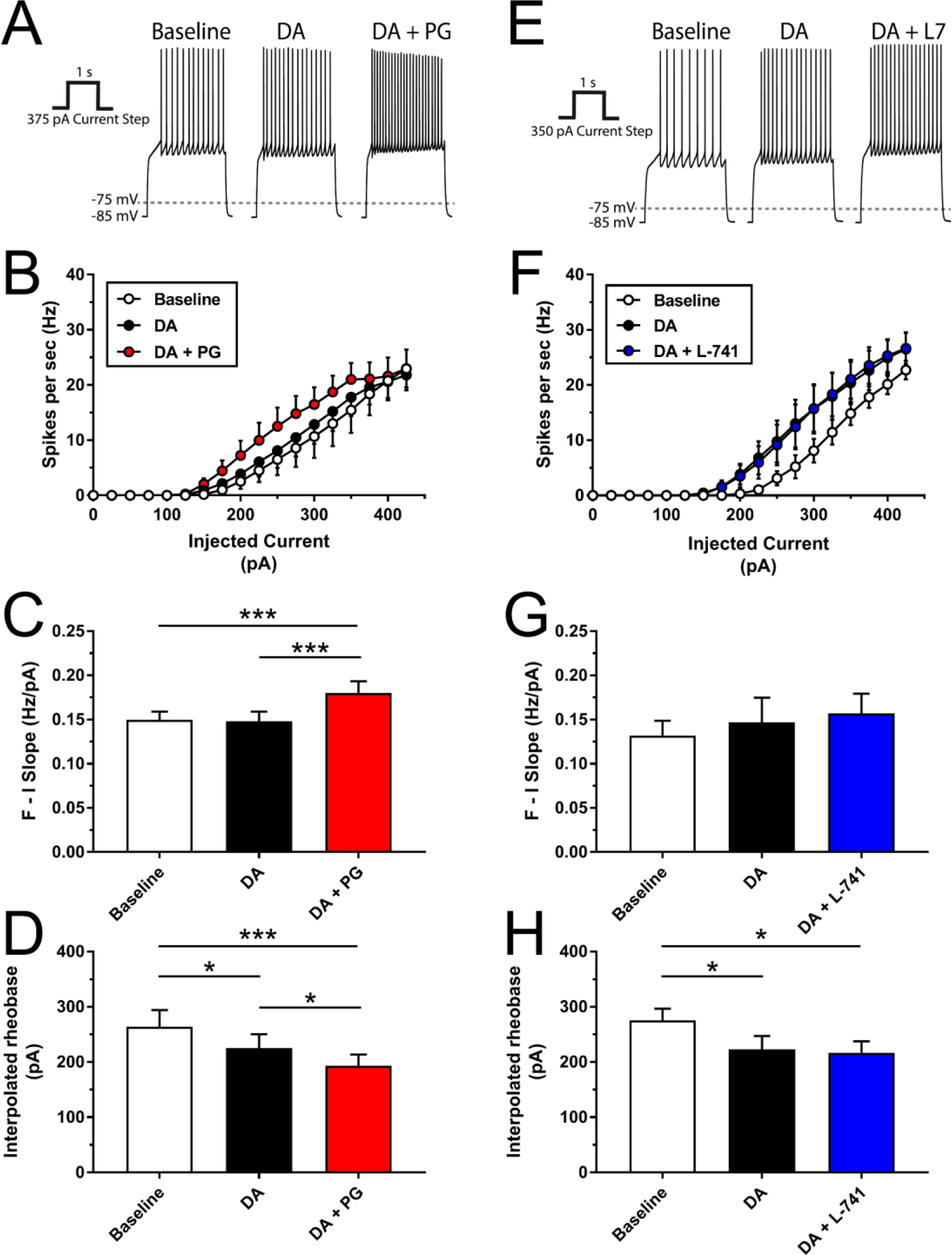
Effects of selective D_2_R or D_3_R antagonism on neuronal activity in D1-MSNs as measured by *ex vivo* slice electrophysiology. **(A)** Representative traces depict the change in firing rate of individual neurons after application of 60 μM DA followed by 60 μM DA + 1 nM L-741,626. **(B)** Firing rate as a function of injected current in D1-MSNs (n = 6) following application of 60 μM DA alone or in combination with 1 nM L-741,626 (two-way RM ANOVA: main effect of current [*F*_(12,60)_ = 67.05, *p* < 0.0001], main effect of condition [*F*_(2,10)_ = 2.33, *p* = 0.148], interaction [*F*_(24,120)_ = 1.93, *p* = 0.011]). **(C)** F-I slope, quantified through the primary linear range of the F-I curve for each individual D1-MSN (n = 6), following perfusion of 60 μM DA alone or in combination with 1 nM L-741,626 (one-way RM ANOVA: main effect of treatment condition [*F*_(2,10)_ = 2.60, *p* = 0.124]). **(D)** Interpolated rheobase in D1-MSNs (n = 6) following application of 60μM DA alone or in combination with 1 nM L-741,626 (one-way RM ANOVA: main effect of treatment condition [*F*_(2,10)_ = 5.69, *p* = 0.022; Fig. 4*D*]). **(E)** Representative traces depict the change in firing rate of individual neurons after application of 60 μM DA followed by 60 μM DA + 1 nM PG01037. **(F)** Firing rate as a function of injected current in D1-MSNs (n = 9) following application of 60 μM DA alone or in combination with 1 nM PG01037 (two-way RM ANOVA: main effect of current [*F*_(12,96)_ = 35.45, *p* < 0.0001], main effect of condition [*F*_(2,16)_ = 7.67, *p* = 0.005], interaction [*F*_(24,192)_ = 0.958, *p* = 0.523]). **(G)** F-I slope, quantified through the primary linear range of the F-I curve for each individual D1-MSN (n = 9), following perfusion of 60 μM DA alone or in combination with 1 nM PG01037 (one-way RM ANOVA: main effect of treatment condition [*F*_(2,16)_ = 13.4, *p* = 0.0004; Fig. 4*G*]). **(H)** Interpolated rheobase in D1-MSNs (n = 9) following application of 60μM DA alone or in combination with 1 nM PG01037 (one-way RM ANOVA: main effect of treatment condition [*F*_(2,16)_ = 12.74, *p* = 0.002]). Data points are depicted as the mean ± SEM. \**p* < 0.05, \*\*\**p* < 0.001, significant difference by Holm-Sidak’s test

The effects of L-741,626 on D2-MSN spike frequency are shown in Figure 5A-D. Application of either DA alone or DA + L-741,626 significantly reduced spike frequency as compared to baseline, but the effects of combined administration of DA + L-741,626 did not significantly differ from the DA alone condition (Figure 5B). None of the treatments significantly affected the slope of the F-I curve as compared to baseline (Figure 5C). In agreement with results on firing rate, rheobase values were significantly increased in the DA alone and DA + L-741,626 conditions as compared to baseline, but did not significantly differ from each other (Figure 5D). We next tested the impact of PG01037 application on D2-MSN spike frequency (Figure 5E-H). As compared to baseline, the application of either DA alone or DA + PG01037 reduced spike frequency (Figure 5F), but the DA alone and DA + PG01037 conditions did not significantly differ from each other. F-I slope was unaffected by any treatment compared to baseline (Figure 5G). Finally, application of DA alone or DA + PG01037 significantly increased rheobase values compared to baseline, but not differ from each other (Figure 5H).

**Figure 5.**
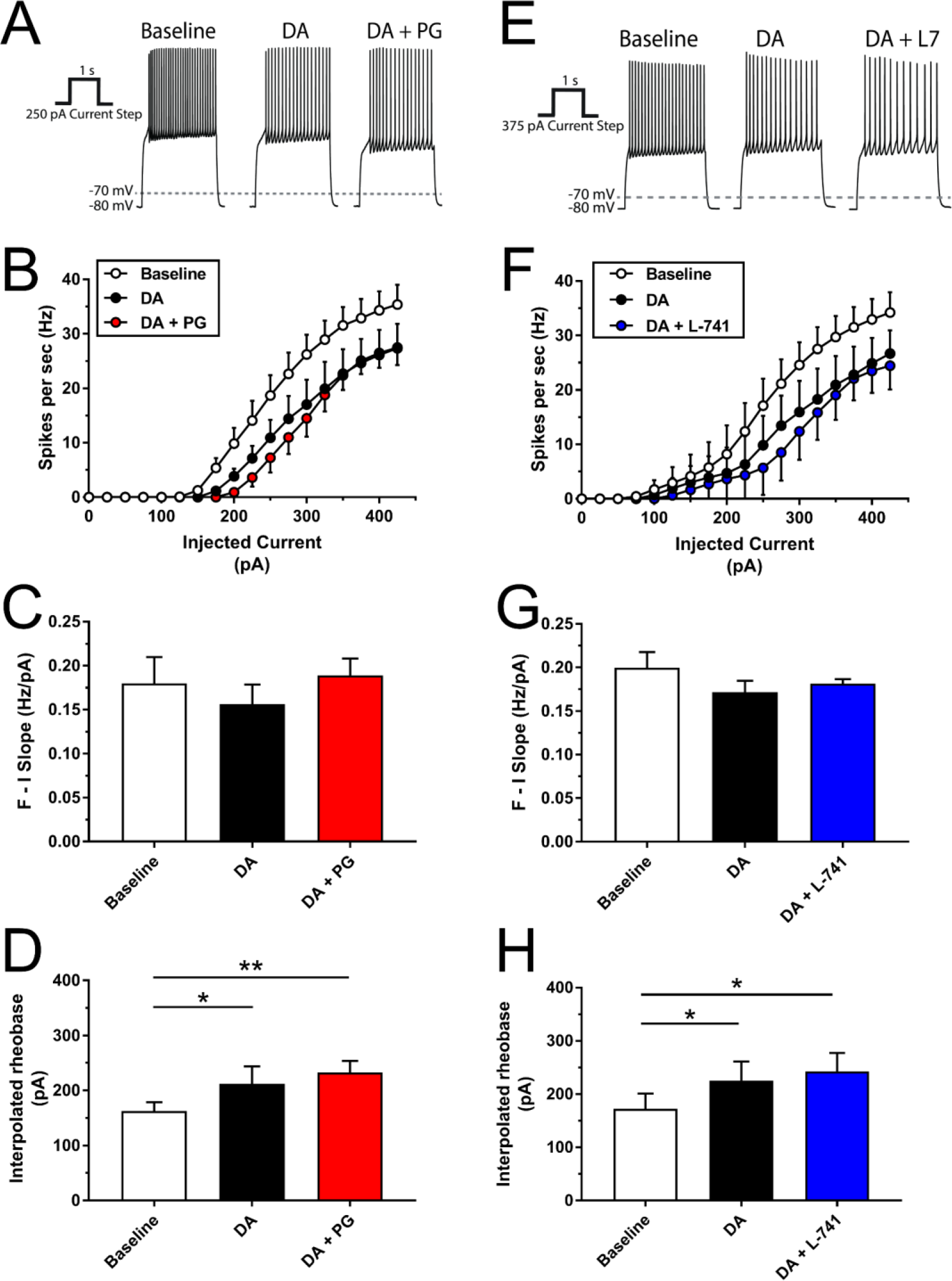
Effects of selective D_2_R or D_3_R antagonism on neuronal activity in D2-MSNs as measured by *ex vivo* slice electrophysiology. **(A)** Representative traces depict the change in firing rate of individual neurons after application of 60 μM DA followed by 60 μM DA + 1 nM L-741,626. **(B)** Firing rate as a function of injected current in D2-MSNs (n = 5) following application of 60 μM DA alone or in combination with 1 nM L-741,626 (two-way RM ANOVA: main effect of current [*F*_(12,48)_ = 60.99, *p* < 0.0001], main effect of condition [*F*_(2,8)_ = 12.43, *p* = 0.004], interaction [*F*_(24,96)_ = 2.65, *p* = 0.0004]). **(C)** F-I slope, quantified through the primary linear range of the F-I curve for each individual D2-MSN (n = 5), following perfusion of 60 μM DA alone or in combination with 1 nM L-741,626 (one-way RM ANOVA: main effect of treatment condition [*F*_(2,8)_ = 2.63, *p* = 0.132]). **(D)** Interpolated rheobase in D2-MSNs (n = 5) following application of 60μM DA alone or in combination with 1 nM L-741,626 (one-way RM ANOVA: main effect of treatment condition [*F*_(2,8)_ = 7.22, *p* = 0.016]). **(E)** Representative traces depict the change in firing rate of individual neurons after application of 60 μM DA followed by 60 μM DA + 1 nM PG01037. **(F)** Firing rate as a function of injected current in D2-MSNs (n = 5) following application of 60 μM DA alone or in combination with 1 nM PG01037 (two-way RM ANOVA: main effect of current [*F*_(12,48)_ = 53.11, *p* < 0.0001], main effect of condition [*F*_(2,8)_ = 13.85, *p* = 0.003], interaction [*F*_(24,96)_ = 4.01, *p* < 0.0001]). **(G)** F-I slope, quantified through the primary linear range of the F-I curve for each individual D2-MSN (n = 5), following perfusion of 60 μM DA alone or in combination with 1 nM PG01037 (one-way RM ANOVA: main effect of treatment condition [*F*_(2,8)_ = 0.82, *p* = 0.478]). **(H)** Interpolated rheobase in D2-MSNs (n = 5) following application of 60μM DA alone or in combination with 1 nM PG01037 (one-way RM ANOVA: main effect of treatment condition [*F*_(2,8)_ = 13.79, *p* = 0.003]). Data points are depicted as the mean ± SEM. \**p* < 0.05, \*\**p* < 0.01, significant difference by Holm-Sidak’s test.

The results obtained from recordings in D2-MSNs indicated that DA reduced the excitability of these neurons, but selective blockade of either D_2_Rs or D_3_Rs alone was incapable of reversing this effect. Because D2-MSNs co-express D_2_Rs and D_3_Rs, we speculated that singular pharmacological blockade of either receptor alone fails to alleviate DA-mediated inhibition because DA binding at the spared receptor subtype is sufficient to exert efficacious inhibitory action on the cell. To test this hypothesis, we first assessed DA-mediated inhibition of spike frequency in D2-MSNs following administration of the nonselective D_2_R/D_3_R antagonist sulpiride. DA alone again produced an expected reduction in spike frequency, but the addition of sulpiride completely abolished this effect (Supplemental Figure S4A-B). We next tested whether co-administration of both L-741,626 and PG01037 would recapitulate the effects of sulpiride. DA alone produced the expected reduction in spike frequency, and simultaneous co-administration of 1 nM L-741,626 + 1 nM PG01037 attenuated this effect (Supplemental Figure S4C-D). These findings suggest that DA-mediated inhibition of D2-MSN activity involves DA binding at both D_2_Rs and D_3_Rs, and that simultaneous blockade of both receptor subtypes may be necessary to temper the impact of DA on the D2-MSN’s firing properties.

### The Enhancement of Cocaine’s Locomotor Stimulant Effects by D_3_R Antagonism Requires Intact D_1_R Signaling

Based on the results of our FSCV and electrophysiological experiments, we hypothesized that D_3_R antagonism potentiates cocaine-induced locomotion in mice via a combination of two distinct neuropharmacological mechanisms in the NAc. First, D_3_R antagonism enhances cocaine-induced increases in DA release, likely via blockade of presynaptic D_3_Rs on DA terminals. Second, D_3_R antagonism results in potentiated excitation of D1-MSNs by DA, likely via blockade of postsynaptic D_3_Rs that normally provide inhibitory tone on these cells. These combined effects of D_3_R blockade create a scenario in which cocaine increases DA levels beyond its normal capacity, and this DA in turn hyperexcites D1-MSNs, thereby producing greater levels of locomotion as compared to cocaine alone. An important facet of this system is that although D_1_R signaling and D_3_R signaling oppose each other in D1-MSNs, the presence of some basal level of D_1_R-mediated excitation within these cells is ultimately required for the behavioral effects of cocaine to emerge. We therefore reasoned that a low level of D_1_R blockade should reduce cocaine-induced locomotion, and that this effect could be reversed with combined treatment of a D_3_R antagonist. However, if the level of D_1_R blockade was substantially strengthened, the effects of a D_3_R antagonist should be rendered ineffective because its capacity to enhance cocaine’s behavioral effects is dependent upon some threshold amount of intact D_1_R signaling. To test these hypotheses, we performed a final set of locomotor activity studies in mice. Pretreatment with a low dose (0.03 mg/kg) of the D_1_-like receptor antagonist SCH23390 caused a modest ~45% attenuation of the locomotor-activating effects of 3.0 mg/kg cocaine, and this effect was reversed by co-administration of 10.0 mg/kg PG01037 (Figure 6A; time course, Supplemental Figure S5A). Increasing the dose of SCH23390 10-fold caused a greater (~63%) reduction in cocaine-induced locomotion, but co-administration of 10.0 mg/kg PG01037 had no impact on the effects of this higher dose of SCH23390 (Figure 6B; time course, Supplemental Figure S5B). Collectively, these findings support the notion that D_3_R antagonism potentiates the stimulant effects of cocaine via a D_1_-like receptor-dependent mechanism.

**Figure 6.**
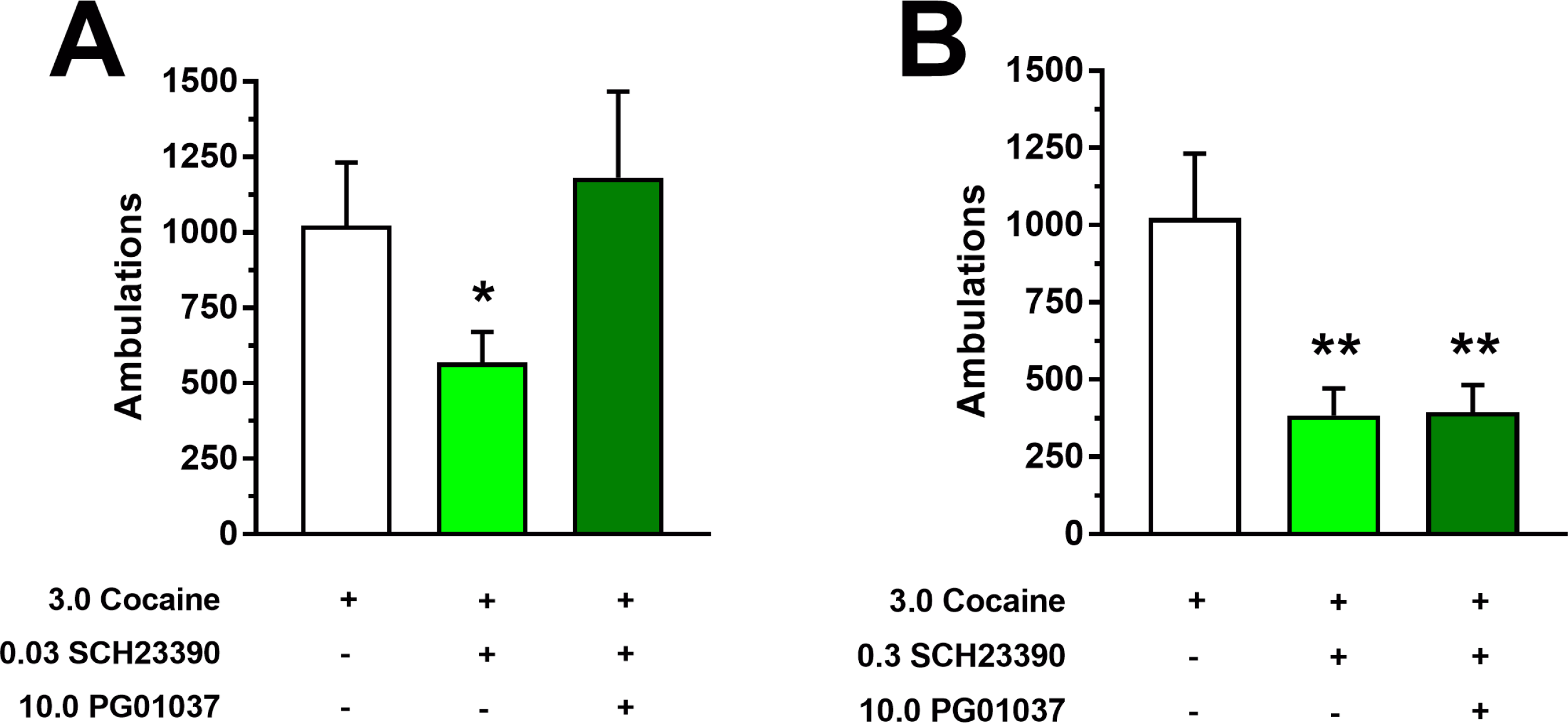
Effects of combined pretreatment with the D_3_R antagonist PG01037 and the D_1_-like receptor antagonist SCH23390 on acute cocaine-induced locomotor activity in mice. **(A)** Total number of ambulations in the 60 min period following administration of 3.0 mg/kg cocaine. 30 min prior to cocaine administration, animals had been injected with a combination of PG01037 (vehicle or 10.0 mg/kg) and SCH23390 (vehicle or 0.03 mg/kg SCH23390). One-way RM ANOVA: main effect of treatment condition, [*F*_(2,14)_ = 6.49, *p* = 0.010]. **(B)** Conditions were identical to panel A, with the exception that the dose of SCH23390 was increased ten-fold (vehicle or 0.3 mg/kg). One-way RM ANOVA: main effect of treatment condition, [*F*_(2,14)_ = 11.78, *p* = 0.001]. Each data point represents mean ± SEM ambulations. All mice received all treatments (*n* = 8). Drug treatments are depicted in a table below each figure; a plus symbol indicates the active dose of the drug was administered, while a minus symbol indicates that its vehicle was administered. \**p* < 0.05, \*\**p* < 0.01, significant difference by Dunnett’s test compared to vehicle/vehicle/3.0 cocaine condition (empty bar)

## Discussion

In the present study, we demonstrate that selective D_3_R antagonism enhances several facets of NAc DA neurotransmission and potentiates the behavioral-stimulant effects of cocaine, while selective D_2_R antagonism produces the opposite behavioral valence and attenuates cocaine’s psychostimulant effect. Furthermore, we identify hyperexcitation of D1-MSNs as a likely mechanism by which D_3_R blockade uniquely facilitates NAc DA neurotransmission and associated behavior.

While nonselective antagonism of D_2_-like receptors has previously been reported to attenuate the locomotor-activating effects of cocaine (68, 69, 72), the specific contributions of the D_2_-like receptor subtypes in mediating these effects have remained unclear. Results from our behavioral studies show that systemic pretreatment with the selective D_2_R antagonist L-741,626 attenuates the acute locomotor-stimulant effects of cocaine and abolishes the development of locomotor sensitization, while administration of the selective D_3_R antagonist PG01037 enhances cocaine’s acute locomotor-stimulant effects and renders a low, ineffective dose of cocaine capable of inducing behavioral sensitization. Intra-NAc infusion of PG01037 recapitulated the effects of systemic administration, thus confirming the NAc as a key neuroanatomical substrate for modulation of cocaine-induced locomotion by D_3_R antagonism. The effects of D_3_R blockade are therefore in direct opposition to those reported previously for nonselective D_2_-like receptor antagonism (68, 69, 72) and for selective D_2_R antagonism in the present study, suggesting that the effect of nonselective D_2_-like receptor blockade on cocaine-induced locomotion is mediated predominantly by antagonism of the D_2_R in the NAc. This is supported by evidence that genetic deletion of the D_2_R subtype within striatal MSNs in mice diminishes the locomotor response to cocaine (73). Moreover, our present results with PG01037 are in agreement with the previous finding that administration of another highly-selective D_3_R antagonist, NGB 2904, potentiates amphetamine-induced locomotor activity in mice (51), as well as the reported hypersensitivity of various lines of D_3_R knockout mice to the behavioral-stimulant effects of cocaine (52, 74) and amphetamine (50, 52). Some studies have failed to detect enhanced stimulant-induced locomotor activity following D_3_R antagonism in rodents, but this discrepancy could be attributed to several factors including the use of compounds with low selectivity for D_3_R vs. D_2_R, use of a limited dose range of psychostimulants, etc. Our within-subjects, multi-dose approach represents a significant advancement over previous investigations and reveals for the first time a clear leftward/upward shift of the cocaine dose-response function, interpreted unequivocally as a pharmacological potentiation. Our observed enhancement of acute and repeated cocaine-induced locomotion following PG01037 administration can be attributed to changes in NAc DA neurotransmission exclusively for several reasons. First, psychostimulant-induced increases in rodent locomotion are strongly linked to the modulation of DA levels specifically within the NAc (75-77). Second, the D_3_R is enriched in the mouse NAc as compared to other regions (23-25). Finally, we show in the present study that intra-NAc infusion of PG01037 is sufficient to produce the enhanced behavioral response to cocaine.

D_2_Rs and D_3_Rs within the NAc are expressed both presynaptically on DAergic terminals and postsynaptically on MSNs (8, 9). Because blockade of presynaptic DA autoreceptors typically results in enhanced presynaptic DA release (78-81), it seemed plausible that D_3_R antagonism may potentiate cocaine-induced locomotion, at least in part, by exacerbating cocaine-induced increases in extracellular DA. In support of this hypothesis, we found that application of PG01037 enhanced cocaine-induced increases in DA release and also significantly facilitated cocaine’s inhibitory action on DA clearance. The effects we observed with PG01037 in our FSCV experiments are similar to those produced by another highly-selective D_3_R antagonist, SB-277011-A, (41, 44), and mirror the facilitation of stimulant-induced increases in extracellular DA following systemic D_3_R antagonism (43). Together, these studies and our present results indicate that presynaptic D_3_R antagonism potentiates stimulant-induced increases in NAc DA levels, even in the absence of any effects on basal DA concentrations. The exact mechanisms by which D_3_R antagonists exert these effects on DA release and clearance and why they only do so under conditions of increased DA levels remain to be determined, but possible explanations include disinhibition of the presynaptic terminal via autoreceptor blockade (8, 26, 27) and/or modulation of the structure and/or function of the dopamine transporter itself (41, 82). Regardless, these actions are unlikely to wholly explain the facilitatory effects of D_3_R antagonism on cocaine-induced locomotion because application of L-741,626 to NAc slices produced effects similar to, and not opposite from, those produced by PG01037. The opposing influences of selective D_2_R and D_3_R antagonism on behavioral output related to NAc DA signaling do not therefore appear to be mediated by disparate effects on presynaptic DA release.

To the best of our knowledge, the present study represents the first assessment of selective D_2_R or D_3_R antagonism on DA-induced changes in D1-MSN and D2-MSN activity within the NAc. With respect to D2-MSNs, although neither L-741,626 nor PG01037 application alone blocked DA-mediated inhibition of D2-MSN activity, we found that their combined administration produced a partial blockade of DA’s effects that resembled the effects of the nonselective D_2_/D_3_ receptor antagonist sulpiride, suggesting that simultaneous D_2_R and D_3_R antagonism may be required to fully prevent the inhibitory actions of DA on these cells. The finding that ineffective doses, when combined, produced a measurable change suggests that these compounds exert qualitatively similar rather than dissimilar effects on D2-MSN activity and that their combined effects are at least additive, if not synergistic, in nature in these cells. In contrast to their effects on D2-MSNs, we observed qualitative differences between selective D_2_R and D_3_R antagonism on DA-mediated excitation of D1-MSNs. The finding that L-741,626 failed to significantly alter the excitatory impact of DA on D1-MSNs was not surprising given the rarity of D_2_R co-expression in these cells (83). However, a significant proportion of D1-MSNs in the NAc co-express the D_3_R (30, 33-35), and accordingly we found that blockade of D_3_Rs via application of PG01037 modulated the activity of D1-MSNs, rendering them hyperexcitable to DA application. D_1_-like receptors are positively coupled to adenylyl cyclase via Gα_s_ protein activation and typically produce intracellular effects that promote neuronal excitation, while the D_3_R is negatively coupled to adenylyl cyclase via Gα_i/o_ protein activation and causes neuronal inhibition (8, 9). When co-expressed in the same cells, these receptors may exert opposing influences on neuronal activity (84). Therefore, the hypersensitivity to DA-mediated excitation exhibited by D1-MSNs in the presence of PG01037 likely reflects the removal of tonic inhibitory tone that would otherwise be provided by DA’s activation of the D_3_R. As D_1_-like receptor activation has been positively linked with stimulant-induced locomotor output (85-88), our findings provide a plausible mechanistic explanation as to how D_3_R antagonism results in exacerbated stimulant-induced locomotion, i.e. through increased activation of NAc D1-MSNs via reductions in D_3_R-mediated inhibition. Consistent with this model, we found that D_3_R antagonism functionally opposed the reductions in cocaine-induced locomotion exerted by a low dose of the D_1_-like receptor antagonist SCH23390, likely because some residual level of D_1_-like receptor signaling remained that was sufficient to engender increased locomotor output when tonic D_3_R-mediated inhibition was antagonized. However, when the dose of SCH23390 was increased ten-fold, the D_3_R antagonist was rendered ineffective, probably because the level of D_1_-like receptor signaling was no longer sufficient to increase locomotor output. Our electrophysiological and behavioral studies are uniformly consistent with the hypothesis that D_3_R antagonism renders NAc D1-MSNs hyperresponsive to DA-induced excitation, and also provide mechanistic evidence as to how D_3_R antagonism functionally enhances the behavioral, cellular, and neurochemical effects of cocaine observed herein and previously by others (38, 41, 44, 74, 89, 90).

In summary, we report that pretreatment with the selective D_3_R antagonist PG01037 enhances, while the selective D_2_R antagonist L-741,626 attenuates, the behavioral-stimulant effects of cocaine in mice. We also demonstrate that the effects of PG01037 are likely mediated by a two-pronged neuropharmacological modulation of NAc DA neurotransmission. First, D_3_R antagonism potentiates cocaine-induced increases in presynaptic DA release, and second, it causes hypersensitivity of NAc D1-MSNs to the excitatory effects of DA while lacking appreciable effects on D2-MSNs. Our results help to resolve a complicated and mixed literature regarding the impact of D_3_R antagonism on NAc-related signaling and behavioral output that will be of importance to those exploring D_3_R antagonists as potential pharmacotherapeutics for various DA-related disorders.

## Acknowledgements

This work was supported by the following funding sources: National Institutes of Health grants K99DA039991 to D.F.M., T32ES012870 to R.C.B., F31DA044726 to S.L.F., F32NS098615 to K.P.S, F31DA037652 and F32AG058396 to K.A.S., ZIADA000424 to A.H.N., R01ES023839 to G.W.M., R21MH113341 and R01MH079276 to C.A.P., R01DA038453 to D.W. and C.A.P., R21DA040788 to D.W.; The Brown Foundation Inc. Fellowship to A.K.P. Portions of this work have previously been reported in poster presentations at the Society for Neuroscience (2016) and the Gordon Research Conference: Catecholamines (2015, 2017).

## References

1. Grace AA (2016): Dysregulation of the dopamine system in the pathophysiology of schizophrenia and depression. Nat Rev Neurosci. 17:524–532.

2. Howes OD, Kapur S (2009): The dopamine hypothesis of schizophrenia: version III--the final common pathway. Schizophr Bull. 35:549–562.

3. Jellinger KA (2014): The pathomechanisms underlying Parkinson’s disease. Expert Rev Neurother. 14:199–215.

4. Vernier P, Moret F, Callier S, Snapyan M, Wersinger C, Sidhu A (2004): The degeneration of dopamine neurons in Parkinson’s disease: insights from embryology and evolution of the mesostriatocortical system. Ann N Y Acad Sci. 1035:231–249.

5. Belujon P, Grace AA (2017): Dopamine System Dysregulation in Major Depressive Disorders. Int J Neuropsychopharmacol. 20:1036–1046.

6. Koob GF, Volkow ND (2016): Neurobiology of addiction: a neurocircuitry analysis. Lancet Psychiatry. 3:760–773.

7. Volkow ND, Morales M (2015): The Brain on Drugs: From Reward to Addiction. Cell. 162:712–725.

8. Beaulieu JM, Gainetdinov RR (2011): The physiology, signaling, and pharmacology of dopamine receptors. Pharmacol Rev. 63:182–217.

9. Missale C, Nash SR, Robinson SW, Jaber M, Caron MG (1998): Dopamine receptors: from structure to function. Physiol Rev. 78:189–225.

10. Newman-Tancredi A, Cussac D, Audinot V, Nicolas JP, De Ceuninck F, Boutin JA, et al. (2002): Differential actions of antiparkinson agents at multiple classes of monoaminergic receptor. II. Agonist and antagonist properties at subtypes of dopamine D(2)-like receptor and alpha(1)/alpha(2)-adrenoceptor. J Pharmacol Exp Ther. 303:805–814.

11. Nordstrom AL, Farde L, Wiesel FA, Forslund K, Pauli S, Halldin C, et al. (1993): Central D2-dopamine receptor occupancy in relation to antipsychotic drug effects: a double-blind PET study of schizophrenic patients. Biol Psychiatry. 33:227–235.

12. Muench J, Hamer AM (2010): Adverse effects of antipsychotic medications. Am Fam Physician. 81:617–622.

13. Miyamoto S, Duncan GE, Marx CE, Lieberman JA (2005): Treatments for schizophrenia: a critical review of pharmacology and mechanisms of action of antipsychotic drugs. Mol Psychiatry. 10:79–104.

14. Wise RA (1995): D1- and D2-Type Contributions to Psychomotor Sensitization and Reward: Implications for Pharmacological Treatment Strategies. Clinical Neuropharmacology. 18.

15. Banks ML, Hutsell BA, Schwienteck KL, Negus SS (2015): Use of Preclinical Drug vs. Food Choice Procedures to Evaluate Candidate Medications for Cocaine Addiction. Curr Treat Options Psychiatry. 2:136–150.

16. Kishi T, Matsuda Y, Iwata N, Correll CU (2013): Antipsychotics for cocaine or psychostimulant dependence: systematic review and meta-analysis of randomized, placebo-controlled trials. J Clin Psychiatry. 74:e1169–1180.

17. Maramai S, Gemma S, Brogi S, Campiani G, Butini S, Stark H, et al. (2016): Dopamine D3 Receptor Antagonists as Potential Therapeutics for the Treatment of Neurological Diseases. Front Neurosci. 10:451.

18. Sokoloff P, Le Foll B (2017): The dopamine D3 receptor, a quarter century later. Eur J Neurosci. 45:2–19.

19. Heidbreder CA, Newman AH (2010): Current perspectives on selective dopamine D(3) receptor antagonists as pharmacotherapeutics for addictions and related disorders. Ann N Y Acad Sci. 1187:4–34.

20. Joyce JN, Millan MJ (2005): Dopamine D3 receptor antagonists as therapeutic agents. Drug Discov Today. 10:917–925.

21. Joyce JN, Millan MJ (2007): Dopamine D3 receptor agonists for protection and repair in Parkinson’s disease. Curr Opin Pharmacol. 7:100–105.

22. Leggio GM, Salomone S, Bucolo C, Platania C, Micale V, Caraci F, et al. (2013): Dopamine D(3) receptor as a new pharmacological target for the treatment of depression. Eur J Pharmacol. 719:25–33.

23. Bouthenet ML, Souil E, Martres MP, Sokoloff P, Giros B, Schwartz JC (1991): Localization of dopamine D3 receptor mRNA in the rat brain using in situ hybridization histochemistry: comparison with dopamine D2 receptor mRNA. Brain Res. 564:203–219.

24. Sokoloff P, Giros B, Martres MP, Bouthenet ML, Schwartz JC (1990): Molecular cloning and characterization of a novel dopamine receptor (D3) as a target for neuroleptics. Nature. 347:146–151.

25. Stanwood GD, Artymyshyn RP, Kung MP, Kung HF, Lucki I, McGonigle P (2000): Quantitative autoradiographic mapping of rat brain dopamine D3 binding with [(125)I]7-OH-PIPAT: evidence for the presence of D3 receptors on dopaminergic and nondopaminergic cell bodies and terminals. J Pharmacol Exp Ther. 295:1223–1231.

26. Chen PC, Lao CL, Chen JC (2009): The D(3) dopamine receptor inhibits dopamine release in PC-12/hD3 cells by autoreceptor signaling via PP-2B, CK1, and Cdk-5. J Neurochem. 110:1180–1190.

27. Diaz J, Pilon C, Le Foll B, Gros C, Triller A, Schwartz JC, et al. (2000): Dopamine D3 receptors expressed by all mesencephalic dopamine neurons. J Neurosci. 20:8677–8684.

28. Fallon JH, Moore RY (1978): Catecholamine innervation of the basal forebrain. IV. Topography of the dopamine projection to the basal forebrain and neostriatum. J Comp Neurol. 180:545–580.

29. Liu XY, Mao LM, Zhang GC, Papasian CJ, Fibuch EE, Lan HX, et al. (2009): Activity-dependent modulation of limbic dopamine D3 receptors by CaMKII. Neuron. 61:425–438.

30. Ridray S, Griffon N, Mignon V, Souil E, Carboni S, Diaz J, et al. (1998): Coexpression of dopamine D1 and D3 receptors in islands of Calleja and shell of nucleus accumbens of the rat: opposite and synergistic functional interactions. Eur J Neurosci. 10:1676–1686.

31. Guitart X, Navarro G, Moreno E, Yano H, Cai NS, Sanchez-Soto M, et al. (2014): Functional selectivity of allosteric interactions within G protein-coupled receptor oligomers: the dopamine D1-D3 receptor heterotetramer. Mol Pharmacol. 86:417–429.

32. Lanza K, Meadows SM, Chambers NE, Nuss E, Deak MM, Ferre S, et al. (2018): Behavioral and cellular dopamine D1 and D3 receptor-mediated synergy: Implications for L-DOPA-induced dyskinesia. Neuropharmacology. 138:304–314.

33. Le Moine C, Bloch B (1996): Expression of the D3 dopamine receptor in peptidergic neurons of the nucleus accumbens: comparison with the D1 and D2 dopamine receptors. Neuroscience. 73:131–143.

34. Schwartz JC, Diaz J, Bordet R, Griffon N, Perachon S, Pilon C, et al. (1998): Functional implications of multiple dopamine receptor subtypes: the D1/D3 receptor coexistence. Brain Res Brain Res Rev. 26:236–242.

35. Surmeier DJ, Song WJ, Yan Z (1996): Coordinated expression of dopamine receptors in neostriatal medium spiny neurons. J Neurosci. 16:6579–6591.

36. Levesque D, Diaz J, Pilon C, Martres MP, Giros B, Souil E, et al. (1992): Identification, characterization, and localization of the dopamine D3 receptor in rat brain using 7-[3H]hydroxy-N,N-di-n-propyl-2-aminotetralin. Proc Natl Acad Sci U S A. 89:8155–8159.

37. Sokoloff P, Diaz J, Le Foll B, Guillin O, Leriche L, Bezard E, et al. (2006): The dopamine D3 receptor: a therapeutic target for the treatment of neuropsychiatric disorders. CNS Neurol Disord Drug Targets. 5:25–43.

38. Congestri F, Formenti F, Sonntag V, Hdou G, Crespi F (2008): Selective D3 Receptor Antagonist SB-277011-A Potentiates the Effect of Cocaine on Extracellular Dopamine in the Nucleus Accumbens: a Dual Core-Shell Voltammetry Study in Anesthetized Rats. Sensors (Basel). 8:6936–6951.

39. Joseph JD, Wang YM, Miles PR, Budygin EA, Picetti R, Gainetdinov RR, et al. (2002): Dopamine autoreceptor regulation of release and uptake in mouse brain slices in the absence of D(3) receptors. Neuroscience. 112:39–49.

40. Koeltzow TE, Xu M, Cooper DC, Hu XT, Tonegawa S, Wolf ME, et al. (1998): Alterations in dopamine release but not dopamine autoreceptor function in dopamine D3 receptor mutant mice. J Neurosci. 18:2231–2238.

41. McGinnis MM, Siciliano CA, Jones SR (2016): Dopamine D3 autoreceptor inhibition enhances cocaine potency at the dopamine transporter. J Neurochem. 138:821–829.

42. Song R, Zhang HY, Li X, Bi GH, Gardner EL, Xi ZX (2012): Increased vulnerability to cocaine in mice lacking dopamine D3 receptors. Proc Natl Acad Sci U S A. 109:17675–17680.

43. Xi ZX, Gardner EL (2007): Pharmacological actions of NGB 2904, a selective dopamine D3 receptor antagonist, in animal models of drug addiction. CNS Drug Rev. 13:240–259.

44. Zapata A, Shippenberg TS (2002): D(3) receptor ligands modulate extracellular dopamine clearance in the nucleus accumbens. J Neurochem. 81:1035–1042.

45. Reavill C, Taylor SG, Wood MD, Ashmeade T, Austin NE, Avenell KY, et al. (2000): Pharmacological actions of a novel, high-affinity, and selective human dopamine D(3) receptor antagonist, SB-277011-A. J Pharmacol Exp Ther. 294:1154–1165.

46. Roberts C, Cummins R, Gnoffo Z, Kew JN (2006): Dopamine D3 receptor modulation of dopamine efflux in the rat nucleus accumbens. Eur J Pharmacol. 534:108–114.

47. Accili D, Fishburn CS, Drago J, Steiner H, Lachowicz JE, Park BH, et al. (1996): A targeted mutation of the D3 dopamine receptor gene is associated with hyperactivity in mice. Proc Natl Acad Sci U S A. 93:1945–1949.

48. Bahi A, Boyer F, Bussard G, Dreyer JL (2005): Silencing dopamine D3-receptors in the nucleus accumbens shell in vivo induces changes in cocaine-induced hyperlocomotion. Eur J Neurosci. 21:3415–3426.

49. Corbin AE, Pugsley TA, Akunne HC, Whetzel SZ, Zoski KT, Georgic LM, et al. (1998): Pharmacological characterization of PD 152255, a novel dimeric benzimidazole dopamine D3 antagonist. Pharmacol Biochem Behav. 59:487–493.

50. McNamara RK, Logue A, Stanford K, Xu M, Zhang J, Richtand NM (2006): Dose-response analysis of locomotor activity and stereotypy in dopamine D3 receptor mutant mice following acute amphetamine. Synapse. 60:399–405.

51. Pritchard LM, Newman AH, McNamara RK, Logue AD, Taylor B, Welge JA, et al. (2007): The dopamine D3 receptor antagonist NGB 2904 increases spontaneous and amphetamine-stimulated locomotion. Pharmacol Biochem Behav. 86:718–726.

52. Xu M, Koeltzow TE, Santiago GT, Moratalla R, Cooper DC, Hu XT, et al. (1997): Dopamine D3 receptor mutant mice exhibit increased behavioral sensitivity to concurrent stimulation of D1 and D2 receptors. Neuron. 19:837–848.

53. Boulay D, Depoortere R, Rostene W, Perrault G, Sanger DJ (1999): Dopamine D3 receptor agonists produce similar decreases in body temperature and locomotor activity in D3 knock-out and wild-type mice. Neuropharmacology. 38:555–565.

54. Xu M, Koeltzow TE, Cooper DC, Tonegawa S, White FJ (1999): Dopamine D3 receptor mutant and wild-type mice exhibit identical responses to putative D3 receptor-selective agonists and antagonists. Synapse. 31:210–215.

55. Karasinska JM, George SR, Cheng R, O’Dowd BF (2005): Deletion of dopamine D1 and D3 receptors differentially affects spontaneous behaviour and cocaine-induced locomotor activity, reward and CREB phosphorylation. Eur J Neurosci. 22:1741–1750.

56. Millan MJ, Dekeyne A, Rivet JM, Dubuffet T, Lavielle G, Brocco M (2000): S33084, a novel, potent, selective, and competitive antagonist at dopamine D(3)-receptors: II. Functional and behavioral profile compared with GR218,231 and L741,626. J Pharmacol Exp Ther. 293:1063–1073.

57. Grundt P, Carlson EE, Cao J, Bennett CJ, McElveen E, Taylor M, et al. (2005): Novel heterocyclic trans olefin analogues of N-{4-[4-(2,3-dichlorophenyl)piperazin-1-yl]butyl}arylcarboxamides as selective probes with high affinity for the dopamine D3 receptor. J Med Chem. 48:839–848.

58. Grundt P, Prevatt KM, Cao J, Taylor M, Floresca CZ, Choi JK, et al. (2007): Heterocyclic analogues of N-(4-(4-(2,3-dichlorophenyl)piperazin-1-yl)butyl)arylcarboxamides with functionalized linking chains as novel dopamine D3 receptor ligands: potential substance abuse therapeutic agents. J Med Chem. 50:4135–4146.

59. Kulagowski JJ, Broughton HB, Curtis NR, Mawer IM, Ridgill MP, Baker R, et al. (1996): 3-((4-(4-Chlorophenyl)piperazin-1-yl)-methyl)-1H-pyrrolo-2,3-b-pyridine: an antagonist with high affinity and selectivity for the human dopamine D4 receptor. J Med Chem. 39:1941–1942.

60. Grundt P, Husband SL, Luedtke RR, Taylor M, Newman AH (2007): Analogues of the dopamine D2 receptor antagonist L741,626: Binding, function, and SAR. Bioorg Med Chem Lett. 17:745–749.

61. Schank JR, Ventura R, Puglisi-Allegra S, Alcaro A, Cole CD, Liles LC, et al. (2006): Dopamine beta-hydroxylase knockout mice have alterations in dopamine signaling and are hypersensitive to cocaine. Neuropsychopharmacology. 31:2221–2230.

62. Weinshenker D, Miller NS, Blizinsky K, Laughlin ML, Palmiter RD (2002): Mice with chronic norepinephrine deficiency resemble amphetamine-sensitized animals. Proc Natl Acad Sci U S A. 99:13873–13877.

63. Kile BM, Walsh PL, McElligott ZA, Bucher ES, Guillot TS, Salahpour A, et al. (2012): Optimizing the Temporal Resolution of Fast-Scan Cyclic Voltammetry. ACS Chem Neurosci. 3:285–292.

64. Lohr KM, Bernstein AI, Stout KA, Dunn AR, Lazo CR, Alter SP, et al. (2014): Increased vesicular monoamine transporter enhances dopamine release and opposes Parkinson disease-related neurodegeneration in vivo. Proc Natl Acad Sci U S A. 111:9977–9982.

65. Hyman SE, Malenka RC, Nestler EJ (2006): Neural mechanisms of addiction: the role of reward-related learning and memory. Annu Rev Neurosci. 29:565–598.

66. Luscher C, Malenka RC (2011): Drug-evoked synaptic plasticity in addiction: from molecular changes to circuit remodeling. Neuron. 69:650–663.

67. Vezina P (2004): Sensitization of midbrain dopamine neuron reactivity and the self-administration of psychomotor stimulant drugs. Neurosci Biobehav Rev. 27:827–839.

68. Baker DA, Khroyan TV, O’Dell LE, Fuchs RA, Neisewander JL (1996): Differential effects of intra-accumbens sulpiride on cocaine-induced locomotion and conditioned place preference. J Pharmacol Exp Ther. 279:392–401.

69. Neisewander JL, O’Dell LE, Redmond JC (1995): Localization of dopamine receptor subtypes occupied by intra-accumbens antagonists that reverse cocaine-induced locomotion. Brain Res. 671:201–212.

70. Aragona BJ, Cleaveland NA, Stuber GD, Day JJ, Carelli RM, Wightman RM (2008): Preferential enhancement of dopamine transmission within the nucleus accumbens shell by cocaine is attributable to a direct increase in phasic dopamine release events. J Neurosci. 28:8821–8831.

71. Hoffman AF, Spivak CE, Lupica CR (2016): Enhanced Dopamine Release by Dopamine Transport Inhibitors Described by a Restricted Diffusion Model and Fast-Scan Cyclic Voltammetry. ACS Chem Neurosci. 7:700–709.

72. Chausmer AL, Katz JL (2001): The role of D2-like dopamine receptors in the locomotor stimulant effects of cocaine in mice. Psychopharmacology (Berl). 155:69–77.

73. Kharkwal G, Radl D, Lewis R, Borrelli E (2016): Dopamine D2 receptors in striatal output neurons enable the psychomotor effects of cocaine. Proc Natl Acad Sci U S A. 113:11609–11614.

74. Carta AR, Gerfen CR, Steiner H (2000): Cocaine effects on gene regulation in the striatum and behavior: increased sensitivity in D3 dopamine receptor-deficient mice. Neuroreport. 11:2395–2399.

75. Delfs JM, Schreiber L, Kelley AE (1990): Microinjection of cocaine into the nucleus accumbens elicits locomotor activation in the rat. J Neurosci. 10:303–310.

76. Kelly PH, Iversen SD (1976): Selective 6OHDA-induced destruction of mesolimbic dopamine neurons: abolition of psychostimulant-induced locomotor activity in rats. Eur J Pharmacol. 40:45–56.

77. Kelly PH, Seviour PW, Iversen SD (1975): Amphetamine and apomorphine responses in the rat following 6-OHDA lesions of the nucleus accumbens septi and corpus striatum. Brain Res. 94:507–522.

78. Benoit-Marand M, Borrelli E, Gonon F (2001): Inhibition of dopamine release via presynaptic D2 receptors: time course and functional characteristics in vivo. J Neurosci. 21:9134–9141.

79. Ford CP (2014): The role of D2-autoreceptors in regulating dopamine neuron activity and transmission. Neuroscience. 282:13–22.

80. Kennedy RT, Jones SR, Wightman RM (1992): Dynamic observation of dopamine autoreceptor effects in rat striatal slices. J Neurochem. 59:449–455.

81. Phillips PE, Hancock PJ, Stamford JA (2002): Time window of autoreceptor-mediated inhibition of limbic and striatal dopamine release. Synapse. 44:15–22.

82. Zapata A, Kivell B, Han Y, Javitch JA, Bolan EA, Kuraguntla D, et al. (2007): Regulation of dopamine transporter function and cell surface expression by D3 dopamine receptors. J Biol Chem. 282:35842–35854.

83. Valjent E, Bertran-Gonzalez J, Herve D, Fisone G, Girault JA (2009): Looking BAC at striatal signaling: cell-specific analysis in new transgenic mice. Trends Neurosci. 32:538–547.

84. Zhang J, Xu M (2006): Opposite regulation of cocaine-induced intracellular signaling and gene expression by dopamine D1 and D3 receptors. Ann N Y Acad Sci. 1074:1–12.

85. Cabib S, Castellano C, Cestari V, Filibeck U, Puglisi-Allegra S (1991): D1 and D2 receptor antagonists differently affect cocaine-induced locomotor hyperactivity in the mouse. Psychopharmacology (Berl). 105:335–339.

86. Le AD, Tomkins D, Higgins G, Quan B, Sellers EM (1997): Effects of 5-HT3, D1 and D2 receptor antagonists on ethanol- and cocaine-induced locomotion. Pharmacol Biochem Behav. 57:325–332.

87. Schindler CW, Carmona GN (2002): Effects of dopamine agonists and antagonists on locomotor activity in male and female rats. Pharmacol Biochem Behav. 72:857–863.

88. Xu M, Hu XT, Cooper DC, Moratalla R, Graybiel AM, White FJ, et al. (1994): Elimination of cocaine-induced hyperactivity and dopamine-mediated neurophysiological effects in dopamine D1 receptor mutant mice. Cell. 79:945–955.

89. Zhang L, Huang L, Lu K, Liu Y, Tu G, Zhu M, et al. (2017): Cocaine-induced synaptic structural modification is differentially regulated by dopamine D1 and D3 receptors-mediated signaling pathways. Addict Biol. 22:1842–1855.

90. Zhang L, Lou D, Jiao H, Zhang D, Wang X, Xia Y, et al. (2004): Cocaine-induced intracellular signaling and gene expression are oppositely regulated by the dopamine D1 and D3 receptors. J Neurosci. 24:3344–3354.

